# Representational confusion: the plausible consequence of demeaning your data

**DOI:** 10.1101/195271

**Authors:** Fernando M. Ramírez

## Abstract

The use of multivariate pattern analysis (MVPA) methods has enjoyed this past decade a rapid increase in popularity among neuroscientists. More recently, similarity-based multivariate methods aiming not only to extract information regarding the class membership of stimuli from their associated brain patterns, say, decode a face from a potato, but to understand the form of the underlying representational structure associated with stimulus dimensions of interest, say, 2D grating or 3D face orientation, have flourished under the name of Representational Similarity Analysis (RSA). However, data-preprocessing steps implemented prior to RSA can significantly change the covariance (and correlation) structure of the data, hence possibly leading to representational confusion—i.e., a researcher inferring that brain area A encodes information according to representational scheme X, and not Y, when the opposite is true. Here, I demonstrate with simulations that time-series demeaning (including z-scoring) can plausibly lead to representational confusion. Further, I expose potential interactions between the effects of data demeaning and how the brain happens to encode information. Finally, I emphasize the importance in the context of similarity analyses of at least occasionally explicitly considering the direction of pattern vectors in multivariate space, rather than focusing exclusively on the relative location of their endpoints. Overall, I expect this article will promote awareness of the impact of data demeaning on inferences regarding representational structure and neural coding.

## Introduction

In the past fifteen years, MVPA methods have enjoyed a rapid increase in popularity within the neuroimaging community^1–3^. Among the goals of early studies applying MVPA was understanding the representational structure of visual percepts^4^, the distribution of object category information in the ventral stream^5–9^, and the hemodynamic correlates of both consciously and unconsciously perceived grating orientation in humans^10,11^. Currently, it is hard to identify a question in cognitive neuroscience that is not being approached with a combination of functional magnetic resonance imaging (fMRI) and MVPA methods.

An interesting recent development in MVPA has been the adoption of classical similarity analysis approaches such as Multidimensional Scaling (MDS,^12–14^) and their adaptation from the domain of psychological variables to that of brain patterns^4,9,15–18^. Thus, while psychologists have traditionally utilized multivariate methods to investigate the structure of mental representational spaces, more recently, neuroscientists have relied on the similarity relationships observed within fMRI, electroencephalographic (EEG) and/or magnetoencephalographic (MEG) patterns, and multichannel intracranial electrophysiological recordings to make inferences regarding representational content and format, as well as claims concerning the tuning properties of neural populations^10,16,17,19–23^. Especially noteworthy, a recent formulation of this similarity-based approach to MVPA termed Representational Similarity Analysis (RSA^18^) was advanced aiming to relate three major branches of systems neuroscience, viz., behavioral measurement, brain measurement, and computational modeling.

In this paper, I use simple simulations to demonstrate that a data pre-processing step which appears to be innocuous in the eyes of some researchers, that is, demeaning of each time series in a dataset consisting of multiple measurement channels, is unsafe when combined with MVPA methods sensitive to the angular relationships among brain pattern vectors—e.g., RSA relying on the linear correlation distance as measure of pattern dissimilarity. These difficulties are mathematically related albeit clearly distinguishable from the unwanted consequences of mean-pattern subtraction in multivariate correlation analyses described by Garrido and colleagues^24^. Moreover, I will argue that because the Euclidean metric is not explicitly sensitive to angular relationships, interpreting pattern analyses based on this dissimilarity measure can be challenging and occasionally even misleading. That is, because this distance measure conflates effects of signal strength with those associated with the direction of a vector in a multidimensional space. Both simulations^21,25,26^ and empirical evidence^27–34^ support the concept that information carried by the direction of pattern vectors, and not their mean, can be specially informative regarding properties of visual objects such as their shape^32^, identity^33^ and three-dimensional (3D) orientation^34^, as well as regarding the form-of-tuning of indirectly sampled neural populations^21,25,26^. Such evidence serves to motivate, at least on these occasions, the use of similarity measures explicitly sensitive to angular relationships among pattern vectors, and, moreover, emphasizes the need to carefully consider the choice of pattern dissimilarity measure in the context of representational similarity analyses.

Critically, I demonstrate that demeaning of multi-channel time series (including z-scoring) prior to MVPA^19,23,35–38^ can significantly change the covariance and correlation structure of the data, and that these changes depend on how information happens to be encoded in a brain area—usually the question under study. Differently said, I exemplify how interactions between the effects of data demeaning and the form of the unknown underlying representational geometry can lead to representational confusion. It may seem evident, on the one hand, that data demeaning can act to mix information associated with signal strength (in fMRI often reflected in regional average responses^39,40^) and that associated with the direction of a pattern vector in multivariate space. Below I show, however, that even in the absence of regional mean differences across conditions, and exclusively due to the form of the underlying representational geometry, data demeaning can lead to representational confusion, namely: inferring that information in brain region A is encoded in accordance with representational scheme X, and not Y, when the opposite is true. Perhaps more gravely, unconventional data pre-processing steps have been invoked^36,37^ to “discard” or “rule out” the influence of global-signal effects which not only cannot discard such effects^25,26^, but can themselves significantly alter the outcome of ensuing MVPA results. Finally, I elaborate on the importance of specifying a meaningful “baseline state” in the context of neuroimaging experiments seeking to make inferences regarding tuning properties of neural populations and representational structure, and discuss strengths and limitations of the currently prevalent approach to estimate a baseline in fMRI studies—i.e., the constant regressor in a standard GLM^41^. Overall, I hope this article will promote awareness regarding the impact of data demeaning on inferences regarding representational structure and neural coding.

## Background

Below, I present seven simulations illustrating possible effects of data demeaning on Representational Similarity Analyses (RSA^18^). In a nutshell, RSA proposes to abstract from brain activity patterns themselves and focus instead on their similarity relationships according to some distance measure (Fig. 1). Thus, if each experimental condition is associated with a brain pattern, an empirical dissimilarity matrix (DSM) can be generated by systematically arranging all pairwise pattern dissimilarities associated with that set of experimental conditions. Such DSMs—or Representational Dissimilarity Matrices (RDMs), if we adopt Kriegeskorte and colleagues’ usage—are believed to roughly encapsulate the information encoded by the neural populations indirectly sampled by fMRI and EEG (Fig. 1*b*). Moreover, these authors propose that several computational models can be related to brain measurements by estimating and comparing the agreement between those models—each expressed as a matrix specifying the expected dissimilarity relationships across conditions given that model—and the empirically estimated DSMs (Fig. 1c). In a similar vein, Kriegeskorte and colleagues suggest relating evidence from different brain measurement techniques by matching the associated DSMs—i.e., abstracting from the patterns themselves and focusing on the corresponding dissimilarity structures. This general approach is attractive, because it circumvents the daunting “correspondency” problem posed by the need to otherwise directly relate model and neural parameters, which may be impossible if a one-to-one mapping between model parameters and data channels cannot be meaningfully established^18^.

In the simulations implemented below, the linear (or Pearson’s) correlation distance (1-*r*) will be used as measure of brain-pattern dissimilarity. That is, because this choice is frequent in the RSA literature, and previously advocated by Kriegeskorte and colleagues^18^. While it must be noted that an alternative distance measure was more recently proposed by Nili and colleagues^42^, namely, crossvalidated linear discriminant t-values (the latter exhibiting interesting and often advantageous properties), nonetheless, to the extent that this distance measure is closely related to the Euclidean distance, it is subject to similar limitations. A key disadvantage is that, by definition, the Euclidean distance conflates vector mean (and more generally speaking, length) and angular effects, thus possibly leading to odd scenarios where neural populations with distinct tuning properties may result in indistinguishable representational distances^21^. In contrast, this would not occur with distance measures explicitly sensitive to angular relationships among vectors, and insensitive to their length—e.g., the cosine and correlation distances. That is, unless distortions on estimates of the true underlying angular relationships were introduced, for example, due to data demeaning (for details, see below).

**Figure 1.**
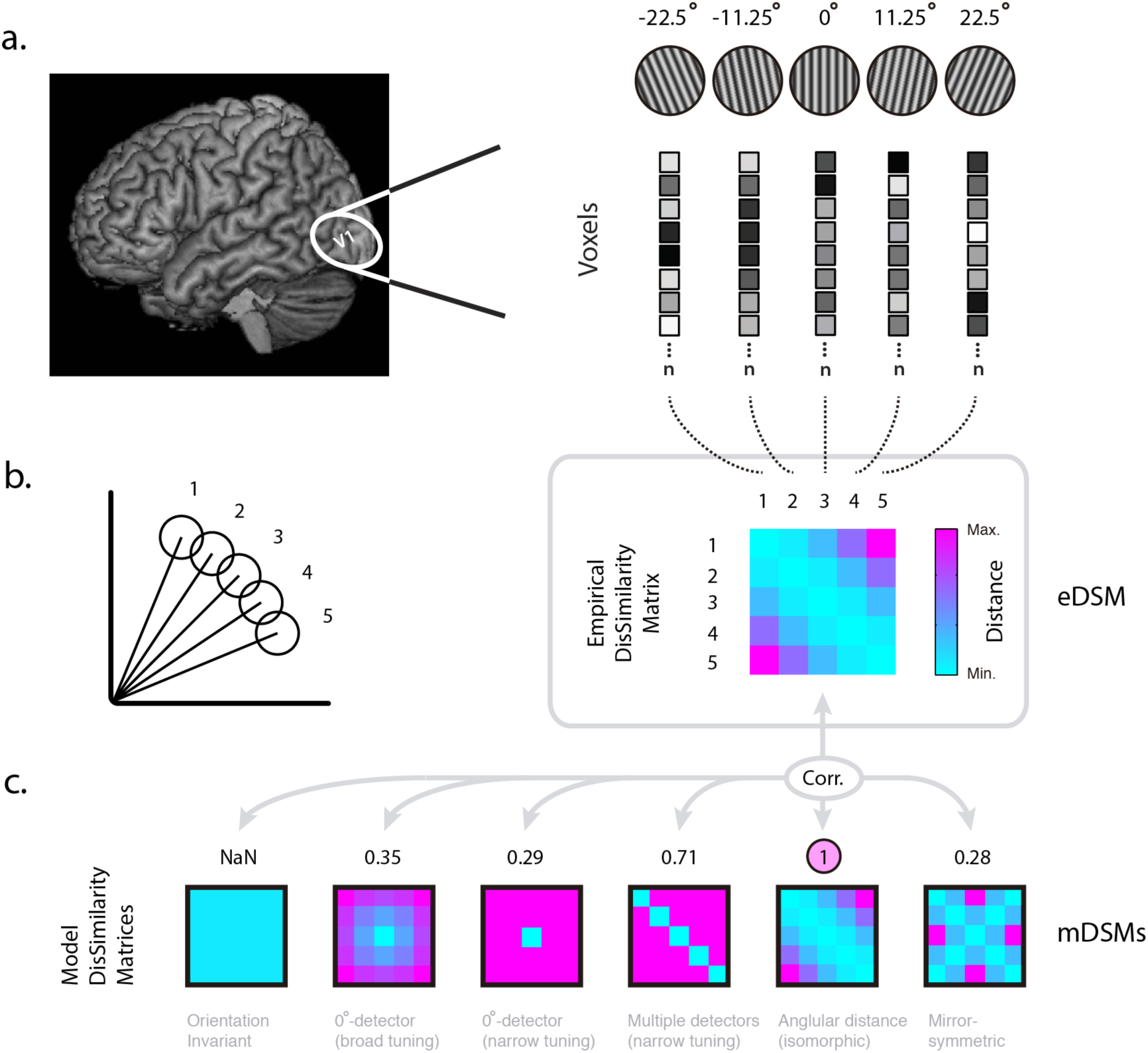
Representational Similarity Analysis: (*a*) Primary visual cortex (V1) is schematically shown within the human brain. A set of activation vectors across V1 is displayed immediately to the right. Each vector depicts the brain response pattern associated with the visual stimulus shown immediately above—i.e., oriented gratings with equal image contrast. (*b*) 2D representation of the *n*-dimensional pattern vectors associated with each of the five oriented gratings shown in *a*. The length of each vector encodes the signal strength associated with each condition. As the angular distance of two gratings increases, so does the associated angular distance in voxel-space. A simulated empirical dissimilarity matrix (eDSM) summarizing all pairwise distances of pattern vectors 1-5 is shown on the right. (*c*) Model dissimilarity matrices associated with each of several candidate representational schemes. Each model DSM is correlated (using Spearman’s rank-order correlation) with the simulated empirical DSM. The model exhibiting the highest correlation—i.e., the best fitting model—is taken to be the most likely underlying representational scheme. In this example, the representational scheme isomorphically reflecting angular disparities among visually presented gratings is the best fitting model.

To remain close to the realm of neurobiological interpretability, the examples and simulations presented below were construed to reflect three imaginary experiments, each assumed to induce effects in primary visual cortex (V1)—where neurons are known to be usually unimodally tuned to a single preferred orientation, and spike-rates positively associated with the observed image contrast levels—or, alternatively, in a fictitious brain area homologous to V1 where neurons respond identically to gratings with orientations equally tilted with respect to the vertical meridian. Let us name this area V1-mirror. Each simulated “experiment” emulates the case were gratings of identical spatial frequency, but variable contrast and orientation, are presented to an observer. Across examples, the contrast levels of the images associated with each orientation are either (I) identical, (II) linearly, or (III) quadratically modulated. Altogether, three experiments, each conducted in V1 and V1-mirror, will serve to specify six examples (see Fig. 2*b*). Please note that neurobiological realism is not of essence here, but only constructing illustrative and conceptually interpretable examples to demonstrate that the rank-order of the entries in the cells of dissimilarity matrices associated with a set of pattern vectors can be strongly changed by data demeaning, and that the associated distortions could lead to representational confusion. Changes in the rank-order of the entries of a dissimilarity matrix (i.e. the ordering of the entries in a matrix after arranging its values in descending order) are key here because (i) dissimilarity matrices are usually taken to reflect the dissimilarity relationships among brain patterns associated with the experimental conditions under investigation, and (ii) similarity in rank-order of empirical and model dissimilarity matrices is the criterion used in RSA for model selection and comparison^18,42^. Thus, rank-order changes due to data transformations could result either in reduced statistical power to detect genuine effects, or—in the worst case—lead to representational confusion. A seventh simulation will serve to illustrate how the distortions demonstrated in examples 1-6 with pattern vectors and cocktail-mean subtraction straightforwardly extend to the case of multi-channel time series demeaning (including z-scoring).

**Figure 2.**
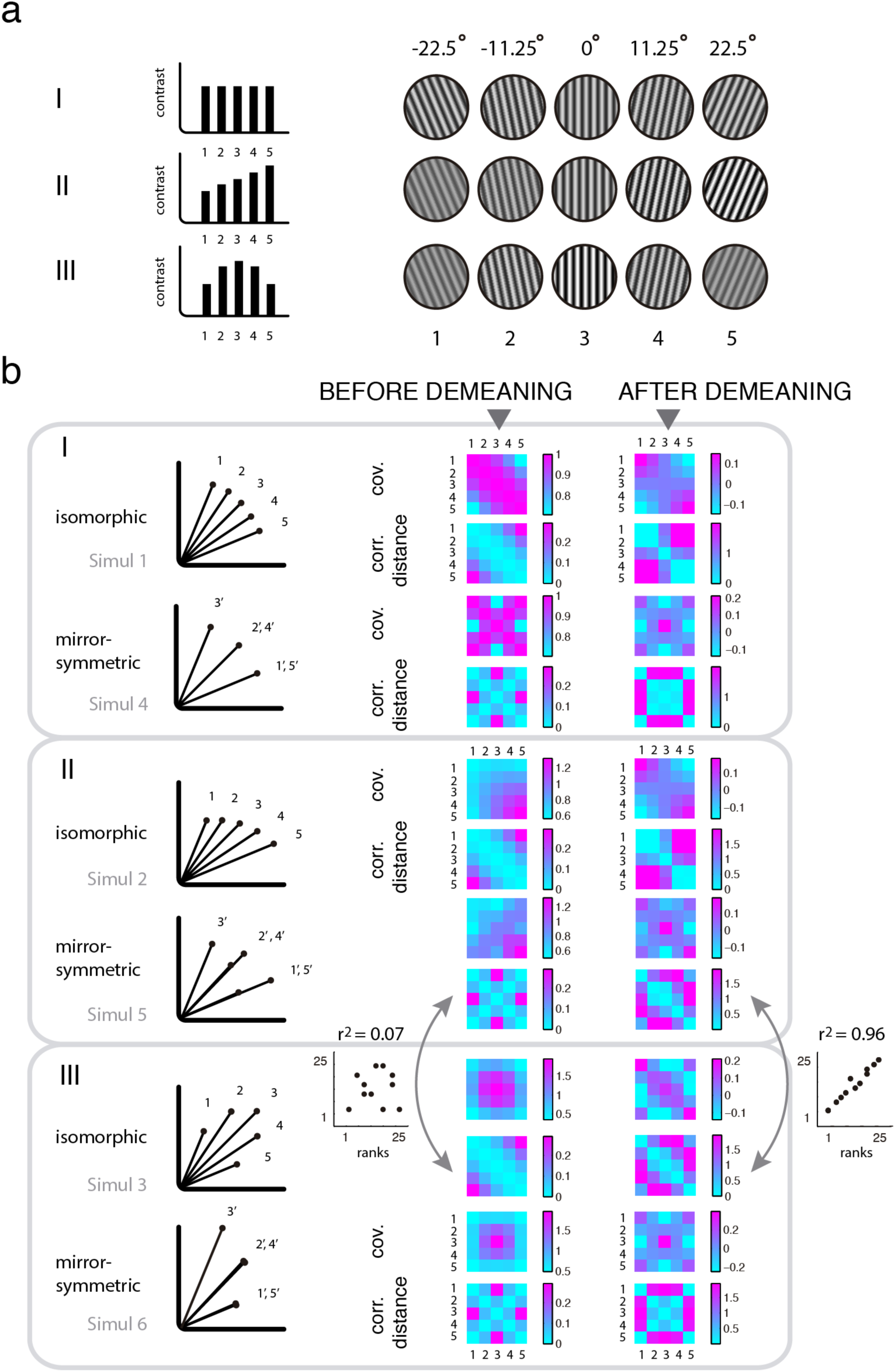
Consequences of cocktail-mean subtraction on pattern dissimilarity structure. (a) Hypothetical visual experiments I-III. Gratings with identical spatial frequency but distinct orientations are shown. In each experiment, image contrast levels are either identical (I), linearly (II), or quadratically modulated (III). (*b*) Hypothetical brain response patters associated with experiments I-III in two distinct brain areas—viz., primary visual cortex (V1) and its envisioned mirror-symmetrically responsive homologue (V1-mirror). Each panel displays brain responses to stimuli shown in I, II, or III. Note that while simulated vectors consist of one-hundred dimensions, they are schematically depicted in two dimensions for illustrative purposes. Vector lengths always reflect grating contrast levels, while angular relationships reflect grating orientations in V1 (top), instead they exhibit identical directions for mirror-symmetrically oriented gratings in V1-mirror (bottom). Each ensemble of 2D-vectors displayed on the leftmost column, the latter used to schematically depict that row’s representational scheme, are accompanied by four matrices. Variance-covariance matrices computed before and after subtraction of the mean pattern across conditions, correspondingly labeled as “before demeaning” and “after demeaning”, are shown on the top row of each sub-panel. Immediately below each pair of covariance matrices, two dissimilarity matrices according to the linear correlation distance are shown, again, before and after demeaning. Changes in the rank-order of both covariance and correlation-distance matrices are evident; compare the appearance of neighboring matrices in each row. Double arrows indicate DSMs before and after demeaning associated with the mirror-symmetric (linearly modulated) and isomorphic (quadratically modulated) encoding schemes (Simulations 5 and 3). Critically, *before* data demeaning these two encoding schemes are easily distinguishable, as indicated by the low correlation observed between the ranks of their associated DSMs (*r*^2^ = 0.07). However, *after* data demeaning these two encoding schemes are hard to distinguish, as indicated by the high correlation observed between the ranks of the associated DSMs (*r*^2^ = 0.96). These observations suffice to demonstrate that cocktail demeaning can introduce changes in representational dissimilarity matrices that could lead to representational confusion.

## Simulations 1-6

Two vectors, each consisting of 100 dimensions (emulating 100 measurement channels), were randomly generated, orthogonalized, and normalized to produce two uncorrelated basis vectors each with a mean of zero and a standard deviation of one. By linearly combining these two basis vectors, two families of vector ensembles were generated:

(1) Isomorphic family: Three vector ensembles were generated, each consisting of five elements (Fig. 2*b* I-III, see top row in each sub-panel). Here, vectors represent brain patterns evoked by visually presented gratings in an observer. Angular disparities among neighboring vectors belonging to this family are 11.25° (π/16 rad) (range = 45°). Thus, angular distances among vectors reflect those observed among the corresponding visually presented gratings^43^. Vector lengths were in each example scaled to reflect the contrast level of the associated gratings (Fig. 2*a*, I-III)—i.e., vector lengths in each group were either identical, linearly, or quadratically modulated, while angular distances remained identical across simulations.

In more detail, a set of vectors *M*_isomorphic_ consisting of five pattern vectors exhibiting the desired angular relationships, zero mean across dimensions, and unit standard deviation, was first obtained by linearly combining 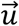 and 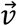 in accordance to the following equation:

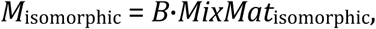

where *B* is a 100- by 2-dimensional matrix comprised by the orthogonal column vectors 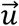 and 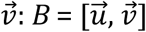. The 2 by 5 dimensional mixing matrix:

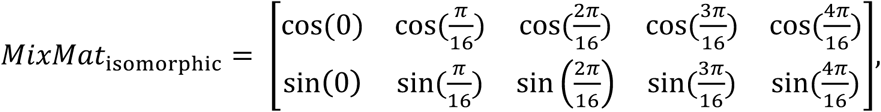

was used to linearly combine the basis vectors in *B* to produce five 100-dimensional vectors, one in each column of *M*_isomorphic_, exhibiting the desired angular relationships, zero means, and unit standard deviations. Then, each column *j* of *M*_isomorphic_ was scaled by multiplication with the correspondingly indexed scalar (*j* = 1, 2, …, 5) in each of the following scaling vectors:

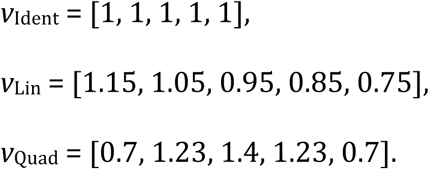

In this way, three ensembles of vectors were generated exhibiting the desired angular relationships, means, lengths, and together constitute the isomorphic family of vectors (see Simulations 1, 2, and 3 in Fig. 2*b*). Please note that the term ‘isomorphic’ is used here only informally to highlight the fact that the angular distances among the orientations of the stimulus gratings are matched by corresponding angular distances among their associated pattern vectors.

(2) Mirror-symmetric family: Three additional ensembles of vectors were generated (Fig. 2*b* I-III, bottom of each panel). In each case, vector 3 represents the pattern associated with the vertically oriented grating. The remaining four vectors exhibit angles proportional to double the *absolute* angular distance between each grating and the vertical meridian. That is, consecutive vectors are separated by angular steps of 22.5° (π/8 rad) (range = 45°). Note that vectors associated with gratings equally removed from the vertical orientation exhibit identical directions, whether the gratings are tilted to the left or to the right.

In more detail, a set of vectors *M*_mirror_ consisting of five pattern vectors exhibiting the desired angular relationships, zero mean, and unit standard deviation was obtained by linearly combining 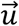 and 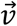 in accordance to the following equation:

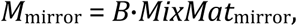

where *B* is the 100- by 2-dimensional matrix comprising the orthogonal basis column vectors 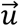 and 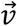 (defined in the previous sub-section). The set of vectors *M*mirror used to generate this second family of vectors was obtained as done above for the isomorphic family but relying on a different mixing matrix, the latter specified as follws,

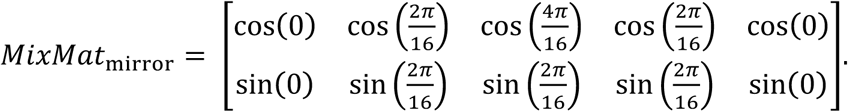

As implemented above for the isomorphic family, the scaling vectors *v*_Ident_, *v*_Quad_, and *v*_Lin_ were used to scale the columns of *M*_mirror_ and hence generate the three ensembles of vectors that constitute this mirror-symmetric family (see Simulations 4-6 in Fig. 2*b*).

### Simulation results (1-6)

By subtracting the mean vector across conditions (i.e. the cocktail mean) from that associated with each condition below I demonstrate that this data transformation can considerably change the rank-order of the entries in the associated covariance and correlation-distance matrices. Compare in Fig. 2*b* the matrices in the middle (before demeaning) and rightmost columns (after demeaning). The fact that neighboring matrices look so strikingly different implies that their rankorder has changed. For instance, this can occur because correlation-distances between pairs of conditions that were among the largest before demeaning can end up among the smallest after demeaning—and vice versa. Furthermore, these examples also show that the correlation-distance matrices associated with two clearly distinct representational schemes before cocktail-mean subtraction—e.g., isomorphic and mirror-symmetric—can become almost indistinguishable after data demeaning. For a dramatic example, compare in Fig. 2*b* the scatterplots of the ranks of the entries of the DSMs indicated with double arrows. Critically, *prior* to data demeaning these two encoding schemes were clearly distinguishable, as indicated by the low correlation observed between the ranks of their associated DSMs (*r*^2^ = 0.07). However, *after* demeaning these two encoding schemes are now hardly distinguishable, as indicated by the high correlation observed between the ranks of the associated DSMs (*r*^2^ = 0.96). In sum, these six examples clearly demonstrate that subtracting the cocktail-mean pattern across conditions can significantly change the angular relationships observed among a set of vectors, alter the rank-order of the entries of the associated covariance and correlation matrices, and therefore possibly lead to representational confusion.

## Simulation 7 (and results)

Figure 3 shows how the effects of cocktail-mean subtraction described immediately above straightforwardly extend to the case of time-series demeaning (including z-scoring). Although this point has seemingly gone unnoticed in the RSA literature, mathematically related artifacts have been discussed in the resting state literature^44–46^. The representational scheme used as ground truth in this seventh simulation is identical to the example isomorphic-III (i.e., with quadratically modulated vector lengths, see Fig. 3*a*), except that here the angular distances between neighboring vectors are 5.625° (π/32 rad) (range 22.5°). Unlike previous simulations, however, in this case one-hundred time series were generated—one per measurement channel. By parametrically manipulating the proportion of non-stimulated periods included in each instance of this simulation, it was possible to investigate the effect of time-series demeaning on RSA inferences as a function of the former parameter. The basic experimental design for the ensuing simulations consists of one single presentation of each oriented grating, interleaved with non-stimulated periods with a duration matching the target proportion of non-stimulated periods specified for each instance of this simulation (see Fig. 3). The signal level specified for all non-stimulated periods is zero. For simplicity, it is further assumed that responses in each simulated measurement channel are perfectly described by zero-lag step functions with a constant duration of 1 min. Clearly, a sound analysis scheme should not lead to inconsistent conclusions regarding representational structure only depending on the proportion of non-stimulated periods included in an experimental design.

**Figure 3.**
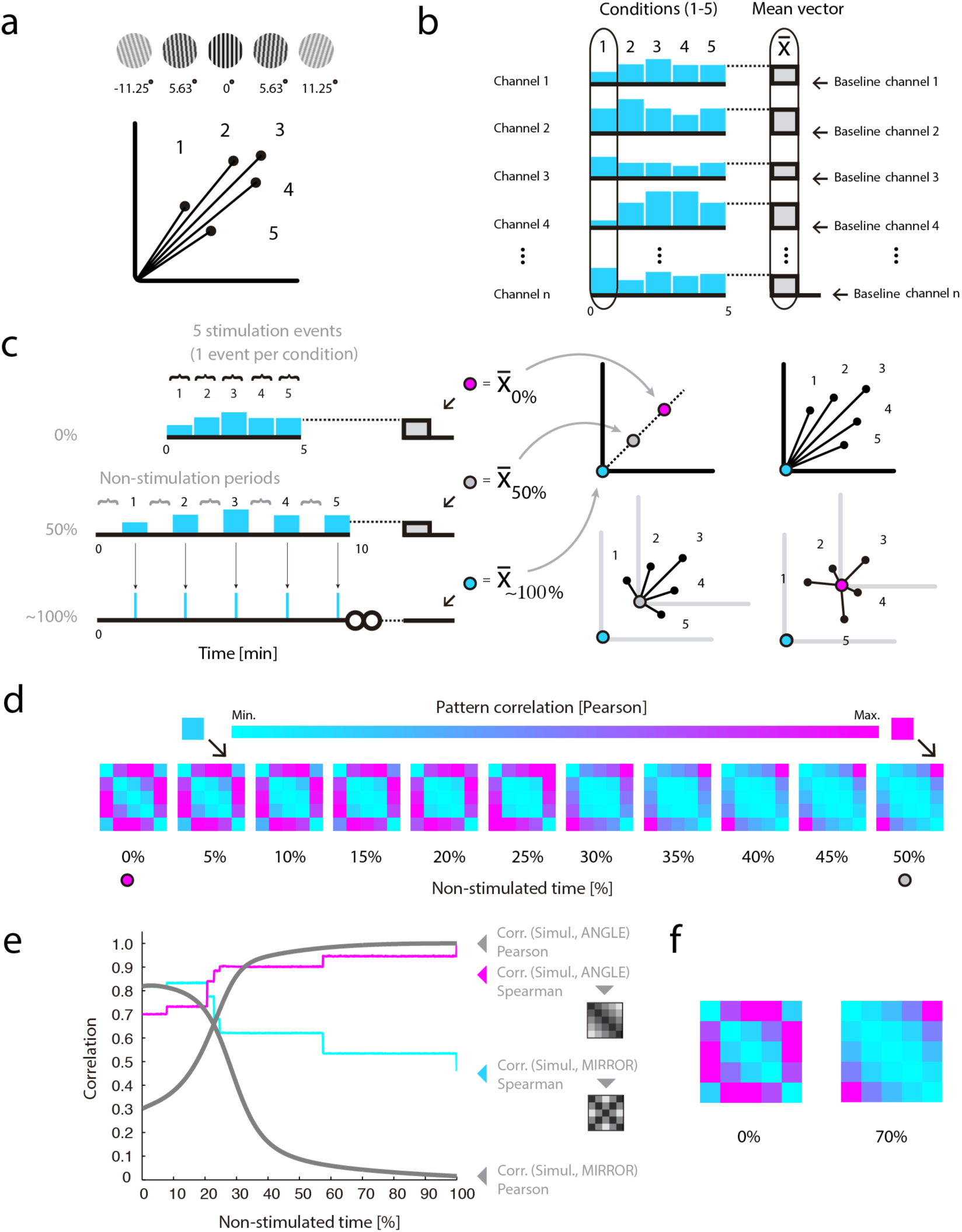
The consequences of cocktail-mean subtraction straightforwardly extend to the case of time-series demeaning. (*a*) Ground-truth for all simulations is a representational scheme were grating angular disparities are isomorphically reflected by the angular disparities between their associated response vectors. Vector lengths reflect image contrast levels. Angular distances between neighboring vectors are 5.625° (range 22.5°). (*b*) Multichannel time series and their mean. In cyan are depicted the signal levels associated with each of five experimental conditions in example measurement channels. On the right, the mean pattern vector across conditions is shown in gray, while outlined in black, also as example, is the pattern associated with condition 1. (*c*) Changing the proportion of pauses in an experimental design influences the mean value observed in each channel. A red dot indicates the mean computed across conditions in one data-channel when the design includes 0% pauses—a setting equivalent here to cocktail-mean subtraction (see text for details). Mean values in designs with 50% and infinite pauses are indicated, respectively, by gray and green dots. On the right, note 2D schematic representations of the location of the multivariate mean associated with designs including different proportions of pauses. The mean pattern varies its location as a function of pause duration, and, critically, on each occasion prescribes distinct angular relationships between the associated pattern vectors. (*d*) Dissimilarity matrices for designs with varying proportions of pauses and always computed after time-series demeaning. The rank-order of the resulting dissimilarity matrices gradually changes as a function of the proportion of non-stimulated periods. Especially noteworthy, arrows indicate a cell that changes from maximal (see DSM for 50% pauses) to almost minimal rank (see DSM for 5% pauses) as a function of the proportion of non-stimulated periods. (*e*) Representational confusion. Shown with separate curves are Pearson and Spearman correlations between the simulated dissimilarity structures (always after demeaning) and two conceptual model templates: (i) a representation monotonically reflecting grating angular disparities (ANGLE), and (ii) a representation mirror-symmetrically encoding grating orientation with respect to the vertical meridian (MIRROR). The results are displayed as a function of the proportion of non-stimulated periods in each simulated experiment. Increasing the proportion of pauses resulted in matrices increasingly similar to the ground truth—here, a representation reflecting angular disparity (see panel *a*). However, note that in the simulations with no or reduced proportions of pauses, both the Pearson and Spearman correlations incorrectly favored the mirror-symmetric model. (*f*) Two matrices—0% and 70% pauses—illustrate significant rank-order differences due to an interaction between the effects of data demeaning and the proportion of pauses in each simulation. Note that while the leftmost dissimilarity matrix exhibits mirror-symmetric features—e.g., note the X-like pattern—the rightmost matrix does not. This interaction of observed dissimilarity structure by proportion of non-stimulated periods could plausibly lead to representational confusion.

Nonetheless, if we envisage the limit case where an experiment includes no pauses, that is, neither before nor after stimulation events, it is easy to see that outcomes arbitrarily close to those due to cocktail-mean subtraction will obtain after time-series demeaning (see Fig. 3*b*). The outcome would be exactly equivalent to cocktail-mean subtraction if neural responses happened to be reflected in our signals without temporal lag and perfectly described by step functions—as conveniently assumed here for didactic purposes. Furthermore, it is key to note that as we consider experiments with increasing proportions of non-stimulated periods, the mean value computed across each time-series will increasingly approximate that of the non-stimulated baseline (set here to a value of zero). In sum, while shorter baseline and inter-stimulus intervals will result in distortions on the associated DSMs increasingly similar to those due to cocktail demeaning, longer pauses will lead to gradually smaller—and eventually no—changes in the rank-order of the resulting DSMs (see Fig. 3*e*). Simulation 7 therefore demonstrates that: (i) time-series demeaning, like cocktail-mean subtraction, can strongly affect ensuing inferences regarding representational structure and neural coding, and (ii) the changes introduced by this data transformation on the rank-order of the ensuing DSMs can strongly depend on arbitrary experimental choices—for instance, the proportion of non-stimulated periods specified by a researcher. That a choice so visibly unrelated to any aspect of the underlying neural populations can so severely influence ensuing inferences regarding neural coding is not only undesirable but arguably also untenable.

Unlike the standard general linear modeling approach (GLM^40^), time-series demeaning and z-scoring do not seek to estimate a baseline level, or to fit a model to the data aiming to minimize the residual error by estimating activation weights for each regressor (conditions) plus an additional constant regressor implicitly modeling the baseline signal level. The demeaning procedures criticized here simply computes the mean across a time series, separately for each measurement channel, and subtracts it from the signal level observed at each time-point. Thus, it must be noted that estimation errors of the baseline level with the GLM approach could in principle also affect the observed rank-order of empirical DSMs and therefore lead to representational confusion. It is easy to see that, as shown to occur with Simulation 7 in the case of multichannel time-series demeaning, inaccurate or biased estimates of the constant (baseline) regressor in a GLM could, for instance, similarly introduce previously inexistent negative correlations between brain patterns. If encountered, such negative correlations may seem hard to explain for a researcher, and, if taken at face value, could lead to erroneous conclusions regarding representational structure and neural coding. In this context, it is worth observing that such distortions, if they occur, are nonetheless expected to be small with the GLM approach compared to those caused by data demeaning—that is, at least in the presence of a roughly orthogonal experimental design and close to canonical hemodynamic response function (HRF).

## The effects of data demeaning interact with the underlying representational geometry

As shown in Figure 2, the changes introduced by data demeaning on the rankorder of empirical DSMs can be strongly influenced by the manner in which the brain happens to encode information—usually the question under study. Thus, given that the true nature of the underlying representational geometry is typically unknown, it follows that the nature of the influences of data demeaning on the empirically observed DSMs will also remain unclear. Moreover, as observed in the previous section, the mean pattern computed across conditions, and therefore the nature of the changes potentially introduced by data demeaning, can be affected by arbitrary characteristics of an experimental design, say, the proportion of non-stimulated periods (see Fig. 3*c*). This is evidently undesirable. For example, if a brain area happened to implement the representational scheme depicted in Fig. 3*a*, then the multivariate mean in an experimental design without pauses would be located on the red dot, with 50% pauses shown in gray, and with close to infinitely long pauses in cyan (Fig. 3*b-c*). Note that after demeaning the pattern-vectors are re-centered and, critically, on each occasion their angular relationships uniquely re-defined. Thus, after data demeaning a researcher may incorrectly interpret the observed dissimilarity structure as evidence that neural populations in V1 exhibit bimodal and mirror-symmetrically tuned response functions (see Fig. 3*d,e*). Moreover, while a researcher investigating V1 with a design including no pauses may conclude that this area encodes information according to a mirror-symmetric scheme, the same researcher relying instead on a design including 50% of non-stimulated periods would reach the opposite conclusion (see Fig. 3*e-f*). In sum, the simulation reported in this section shows that the uncritical use of data demeaning prior to the implementation of RSA can lead to representational confusion, and that the effects of data demeaning can interact with the very form of the unknown underlying representational geometry under investigation. Taken together, these pitfalls speak against the practice of uncritically demeaning your data before conducting RSA analyses.

## Discussion

The simulations presented in this paper serve to demonstrate that it is unsafe to demean your data before conducting MVPA variants (including RSA) sensitive to the angular relationships among estimated response patterns. Because the angular relationships among pattern vectors can be crucially informative regarding the nature of the underlying neural population codes (a concept supported by simulations and empirical data^21,29,33,38,47^), it is evident that at least in those occasions re-centering of time series by subtracting the mean across conditions should be avoided. Moreover, it is also evident that careful attention should be paid when interpreting Euclidean distances if chosen as measure of pattern dissimilarity. One may argue that the Euclidean distance is preferable to the correlation and cosine distances in the context of RSA precisely because it is (at least within runs) unaffected by data re-centering^42,48^. However, I think a researcher’s priority is not to choose a metric insensitive to the possible distortions introduced by data preprocessing steps, but one that faithfully reflects the neural properties under investigation. In my view, an important question in neuroscience is to understand how tuning properties of neural populations sampled by brain measurements may be discovered from their associated multivariate response patterns—thus aiding us to learn from such signals how neurons collectively encode cognitive and sensory variables.

Do my arguments imply that conclusions drawn in previous studies relying on a combination of brain measurements and RSA are necessarily mistaken? No, they do not imply this. However, to the extent that data was demeaned prior to RSA, it can be said that if these conclusions are true, it is not by virtue of the analysis schemes used to provide evidence in their favor. To the extent that studies intend to make claims about neural population tuning properties, the conclusions drawn could change when analyzed in a manner aiming to preserve angular relationships among brain patterns with respect to a pre-stimulation baseline level, thus avoiding re-centering operations that recode brain patterns with respect to an arbitrary origin that seems to me hardly interpretable in neurobiological terms.

Thus, it becomes apparent in the context of MVPA that specifying what constitutes a sound “baseline” level—as well as considering the possible influence on eDSMs of common pattern components^49^—may prove critical when it comes to draw inferences regarding neural population tuning. While some researchers may disagree, I think that the baseline level estimated in the context of GLM analyses can prove to be biologically interpretable, and provide a meaningful reference point to specify angular relationships among brain pattern vectors. More precisely, while in the context of fMRI the estimated baseline value itself may prove hard or even impossible to interpret, the *direction* of the deviations from the estimated baseline evoked by an experimental manipulation may prove to be interpretable in neurobiological terms—e.g., under specific conditions, the direction of an activation pattern may turn out to be uniquely informative regarding the form-of-tuning of indirectly sampled neural populations^21,25,26^. Under this analysis, what matters would be not so much the precise location of the endpoint of a pattern vector in multidimensional space, but how it is that this point actually got there in the first place.

In the context of standard GLM analyses of fMRI experiments, the measured signal levels associated with each condition are usually modeled as activations or deactivations with respect to an implicitly estimated baseline level, resulting in positive or negative weights (parameter estimates, or regression coefficients) associated with each experimental condition. Positive modulations with respect to baseline are probably the most straightforwardly interpretable effects, while negative modulations may require further qualifications—reviewed by Logothetis^50^. Surely, an estimated baseline level could on different occasions reflect physically and cognitively distinguishable states. This implies that the baseline state, as estimated by the standard GLM approach, will on occasions fail to distinguish meaningful brain states. However, this approach seems suited to model event-related modulations in the activity levels of neural populations tuned to specific stimulus features. In my view, it has the further advantage of avoiding haphazard changes in the estimated angular relationships among pattern vectors.

It has been argued that the practice of data demeaning is motivated by the intention to remove from a set of response patterns a presumed shared component^48^. It is key to realize that computing the mean across conditions neither generally nor adequately estimates such putative common component. First, it must be underscored that the concept that such common component even exists is usually nothing more than an assumption. Second, even if such a common component did actually exist, it could in principle (if not in practice) lie in any direction, and exhibit any amplitude. Third, it is often unclear what exactly researchers mean when they speak about such “common component”. If by this they mean that a population of neurons exists in a particular brain area that identically responds to all experimental conditions, then, there is no a priori reason to expect that their associated pattern component should necessarily lie on the location of the cocktail mean. Therefore, unless a plausible justification can be provided explaining why the cocktail mean should accurately estimate a presumed common component (if it exists) this practice should not be uncritically accepted in the context of MVPA analyses sensitive to angular relationships among pattern vectors, including RSA. Nonetheless, it is also important to understand that if a common component were present in the data, and our analyses failed to account for its influence, inferences regarding neural coding based on the RSA methodology could also fail, and hence also lead to representational confusion. This realization serves to highlight the importance of recent efforts to subsume RSA within a more general framework also concerned with encoding and pattern-component models^51^. Likewise, it serves to highlight the fact that if the influence of the measurement process on the observed pattern dissimilarities is not accounted for by such models^21,25,26,52^, they could also lead to representational confusion.

In conclusion, I hope this article will serve as an invitation to reflect on implicit assumptions bearing on the interpretation of MVPA and RSA, and promote awareness of the impact of data demeaning and choice of representational dissimilarity measure on inferences regarding representational structure and neural coding.

## Acknowledgements

First and foremost, I thank Carsten Allefeld for valuable contributions to the ideas advanced in this paper, as well as comments to two previous versions of this manuscript. Furthermore, I thank Lúcia Garrido, Joram Soch, and Sebo Uithol for useful comments to an earlier version of this manuscript, and Kai Görgen for feedback to an early oral presentation of the simulations included in this article.

